# CRISPR-HOLMES-based NAD^+^ detection

**DOI:** 10.1101/2023.12.06.570494

**Authors:** Songkuan Zhuang, Tianshuai Hu, Hongzhong Zhou, Shiping He, Jie Li, Yuehui Zhang, Dayong Gu, Yong Xu, Yijian Chen, Jin Wang

## Abstract

Studies have indicated that intracellular nicotinamide adenine dinucleotide (NAD^+^) level is associated with the occurrence and development of many diseases. While traditional NAD^+^ detection techniques are time-consuming and may require large and expensive instruments. We recently found that the CRISPR-Cas12a protein can be inactivated by AcrVA5-mediated acetylation and reactivated by CobB, using NAD^+^ as the co-factor. Therefore, here in this study, we created a CRISPR-Cas12a-based one-step HOLMES(NAD) system for rapid and convenient NAD^+^ detection with the employment of both acetylated Cas12a and CobB. In HOLMES(NAD), acetylated Cas12a losses its *trans*-cleavage activates and can be reactivated by CobB at the presence of NAD^+^, cutting ssDNA reporters to generate fluorescence signals. HOLMES(NAD) shows both sensitivity and specificity in NAD^+^ detection and can be used for quantitative determination of intracellular NAD^+^ concentrations. Therefore, HOLMES(NAD) not only provides a convenient and rapid approach for target NAD^+^ quantitation but also expands the application scenarios of HOLMES to non-nucleic acid detection.

## 1. Introduction

Nicotinamide adenine dinucleotide (NAD^+^) is essential for cellular catabolic metabolisms, participating as a coenzyme in redox reactions as well as serving as an important co-substrate for NAD^+^-dependent enzymes including sirtuin family of protein deacetylases and the PARP family of DNA repair enzymes. As a result, quantitative detection of the NAD^+^ level may help reflect the status of an organism and preclude the onset and progression of metabolism-related diseases.

So far, a wide variety of technologies have been developed for the measurements of NAD^+^, including high-performance liquid chromatography (HPLC) (Jones 1981), liquid chromatography/electrospray ionization tandem mass spectrometry (MS) (Yamada et al. 2006), enzymatic assays (Bernofsky and Swan 1973) and genetically encoded fluorescent biosensor (Yu et al. 2019). However, these techniques are either time-consuming or may require large and expensive instruments, bringing much inconvenience. For example, the HPLC method has low sensitivity and requires more samples when compared with other techniques. Although the MS method possesses high accuracy and high specificity, it relies on expensive precision instruments and professional operations. The enzymatic method, also known as the colorimetric NAD^+^ assay, needs two steps to calculate the NAD^+^ amount. First, the total amount of NAD^+^ and NADH as well as the amount of NADH are measured, and then the NAD^+^ concentration was indirectly obtained by subtraction. Although the bioluminescent method (Yu et al. 2019) has shown advantages in sensitivity, rapidness and convenience, its specificity need further improvement as it may have difficulty in distinguishing low-concentration NAD^+^ from high-concentration NADH samples. Therefore, it is necessary to develop a rapid, reliable, cost-effective, and quantitative method for convenient NAD^+^ detection.

The clustered regularly interspaced short palindromic repeat (CRISPR)/CRISPR associated (Cas) systems have been found in many bacteria and most archaea, and are well known as an effective defense means for hosts to prevent invasion from mobile genetic element (MGEs), such as bacteriophages (Mojica et al. 2005). Briefly, Cas proteins specifically recognize and cut target DNA under the guidance of guide RNAs, which findings have greatly promoted precise genome editing (Jinek et al. 2012). Among these Cas proteins, Cas12 and Cas13 possess not only *cis*-cleavage activities but also *trans*-cleavage activities, which activities have been employed to develop the next-generation CRISPR diagnostic systems such as Cas12-based HOLMES and Cas13-based SHERLOCK (Gootenberg et al. 2017; Li et al. 2018; Linxian et al. 2019). Previous studies showed that Cas effectors such as Cas12a could be regulated by post-translational acetylation (PTM) of a key lysine site by a GNAT-family acetyltransferase, namely AcrVA5, resulting in its loss of the ability to interact with the protospacer adjacent motif (PAM) site and failure in protecting the hosts by cleaving the invading MGEs (Dong et al. 2019; Knott et al. 2019). Recently, our study demonstrated that AcrVA5-inactivated Cas12a could be reactivated by CobB-mediated deacetylation in a NAD^+^-dependent manner, and reactivated Cas12a recovers its abilities in both target double-stranded DNA (dsDNA) recognition and cleavage *in vitro* and host defense *in vivo* (Kang et al. 2022). Therefore, one could conclude that NAD^+^ plays an important role in regulation of Cas12a activities, including the *trans*-cleavage activities.

Based on above findings, we here employed this mechanism to develop a Cas12a-based CRISPR diagnostic system, namely HOLMES(NAD), for convenient NAD^+^ detection. Our results demonstrate the outstanding performance of HOLMES(NAD), characterized by a low detection limit (22.5 nM), cost-effective, high specificity, and remarkable repeatability (RSD= 3.80-5.82 %) and precision (RSD= 1.65-8.72 %). More importantly, HOLMES(NAD) can quantitatively detect NAD^+^ levels in real biological sample within 30 minutes, following a one-pot and one-step procedure.

## 2. Materials and methods

### 2.1. Construction of plasmids

To construct the LbCas12a expression plasmid pCDF-PylT-LbCas12a, the LbCas12a gene was amplified from plasmid pET28a-LbCas12a (ToloBio) with primers of LbCas12a-F/R with high fidelity DNA polymerase, and the amplicons were purified and digested with BamHI and SalI before being inserted into the same sites in pCDF-PylT (Neumann et al. 2009) *via* T4 DNA ligation.

### 2.2. Purification of protein

Purification procedures of the recombinant proteins were the same as previously described (Kang et al. 2022). Briefly, to obtain acetylated LbCas12a, plasmids of pCDF-PylT-LbCas12a and pET28a-AcrVA5 (Dong et al. 2019) were co-transformed into *E. coli* BL21 (DE3) and the transformants were induced by addition of IPTG at a final concentration of 0.2 mM. Acetylated LCas12a proteins were purified following the steps of Ni-NTA column (HisTap™ HP, GE Healthcare), ion exchange chromatography (HiTrap™ SP HP, GE Healthcare) and molecular sieve (HiLoad™ 16/600, Superdex™ 200 pg, GE Healthcare). Proteins were quantitated using the Bradford method and stored at -80 □ in the storage buffer containing 50 mM Tris-HCl, 600 mM NaCl, 2 mM DTT, 0.2 mM EDTA, 50% Glycerol. The purification of CobB was the same as what we described before (Kang et al. 2022).

### 2.3. Generation of dsDNA target

To create short dsDNA, two complementary oligonucleotides of both forward and reverse were synthesized (Table S2) (Sangon, China) and annealed at a ratio of 5:1 in 1× Taq buffer to form dsDNA, which was then stored at -20 □. The generated dsDNA was diluted to 100 nM with nuclease-free water before use.

To generate FAM-labelled double stranded DNA targets, the target DNA fragment was amplified from plasmid pClone007s-COL1A2 (Linxian et al. 2019) with primers of M13-F and M13-R-FAM with a high-fidelity DNA polymerase. Purified amplicons were then quantitated with NanoDrop 2000 (Thermo Fisher Scientific) and stored at -20 □.

### 2.4. CobB-mediated deacetylation of acLbCas12a

To the perform the deacetylation reaction, 1 μM acLCas12a was treated with 1 μM CobB with the addition of 1 mM NAD^+^ in 1× NEB buffer 3.1 in a 20 μL reaction system. Reaction was carried out at 37 □ for 60 min, and the acetylation status of acLCas12a was then analyzed by western blot.

### 2.5. *cis*- and *trans*-cleavage assays

For *in vitro cis*-cleavage assay, CobB-treated acLCas12a was incubated in a 20 μL reaction system containing 150 ng target dsDNA, 1 μM acetylated LCas12a, 1 μM CobB, 1 μM crRNA, 1 mM NAD^+^. Reaction was performed at 37 □ for 60 min in 1× NEB buffer 3.1, and then stopped by heating at 80 □ for 15 min, followed by immediately chilling on ice before being further analyzed by PAGE.

For *in vitro trans*-cleavage assay, CobB-treated acLCas12a was incubated in a 20 μL reaction system containing 0.5 μM acetylated LCas12a, 1 μM CobB, 0.5 nM target DNA, 12.5 nM crRNA, 0.5 μM 8C FQ-reporter and 0.5 μM NAD^+^. Reaction was performed at 37 □ in 1× NEB buffer 3.1, and results were detected in a real-time PCR machine (Applied Biosystems QuantStudio 3) for 30 min with the fluorescence signal collected every 15 s (λex: 488 nm, λem: 535 nm).

### 2.6. Measurement of cellular NAD^+^ concentration

#### (1) Extraction of cellular NAD^+^

To measure the NAD^+^ levels in cancer cells, cells were first digested with 0.25 % trypsin-EDTA (Gibco), rinsed twice with pre-cooled 1× PBS (HyClone), and then harvested by centrifugation at 5,000 rpm for 5 min at 4 □. Meanwhile, cell numbers were determined by Countless 3 (Thermo Fisher). Cell pellet was resuspended in 300-600 μL acidic extraction buffer (0.3 M HCl) according to the cell numbers, incubated at 95 □ for 5 min and then cooled immediately on ice, followed by centrifugation at 12,000 rpm for 10 min at 4 □. Acidic extraction solution (200 μL) containing NAD^+^ was neutralized using 180-200 μL alkaline buffer (0.3 M KOH) before being stored at -80 □ for the following analysis.

#### (2) Detection of NAD^+^

To detect NAD^+^, 1-2 μL extraction solution containing NAD^+^ were mixed with *trans*-cleavage system containing 0.5 μM acetylated LCas12a, 1 μM CobB, 0.5 nM target DNA, 12.5 nM crRNA and 0.5 μM 8C-FQ-reporter. Reaction was performed at 37 □ in 1× NEB buffer 3.1, and the results were detected in a real-time PCR machine for 30 min with the fluorescence signals collected every 15 s (λex: 488 nm, λem: 535 nm). Finally, the NAD^+^ concentration was calculated according to the standard curve (Y=388.5X + 36523) and the dilution factor.

In parallel, cellular NAD^+^ levels were determined by both MS and the NAD^+^/NADH assay kit with MTT (Cominbio and Solarbio), following the manufacturers’ instructions. For the MTT method, cells were washed with cold PBS, and were then treated with acidic extraction buffer to release NAD^+^ for subsequent determination by reduction of MTT to formazan by NADH, which was generated by NAD^+^ reductive reactions with alcohol dehydrogenase.

### 2.7. Cell culture and pharmacological compounds treatment

Cell lines of HeLa and 97H were maintained in DMEM/high glucose (HyClone) supplemented with 10 % FBS (Gibco) and 1× penicillin-streptomycin (Gibco). Cell lines of HCT116 were maintained in McCoy’s 5A medium (Sigma) supplemented with 10 % FBS (Gibco) and 1× penicillin-streptomycin (Gibco). For pharmacological compounds treatment, cells at ~60 % confluence were pre-incubated with FK886 (Sigma-Aldrich) at a final concentration 100 nM and continually cultured for another 24 h before being counted and collected.

## 3. Result and discussion

### 3.1. Principles of HOLMES(NAD)

Briefly, the HOLMES(NAD) system comprises acetylated Cas12a, CobB, crRNA, target dsDNA and single-stranded DNA (ssDNA) FQ-reporters. As acetylated Cas12a losses *trans*-cleavage activities with target dsDNA as the template, ssDNA FQ-reporters will not be cleaved by acetylated Cas12a, and no fluorescence signal could be detected. Only at the presence of NAD^+^, CobB then removes the acetyl unit from acetylated Cas12a and restores its *trans*-cleavage activities, cleaving the ssDNA FQ-reporters and illuminating fluorescence signals (Fig. 1). Considering the strong relationship between NAD^+^ and the fluorescence signals in HOLMES(NAD), the system can theoretically be employed for NAD^+^ detection.

**Fig 1.**
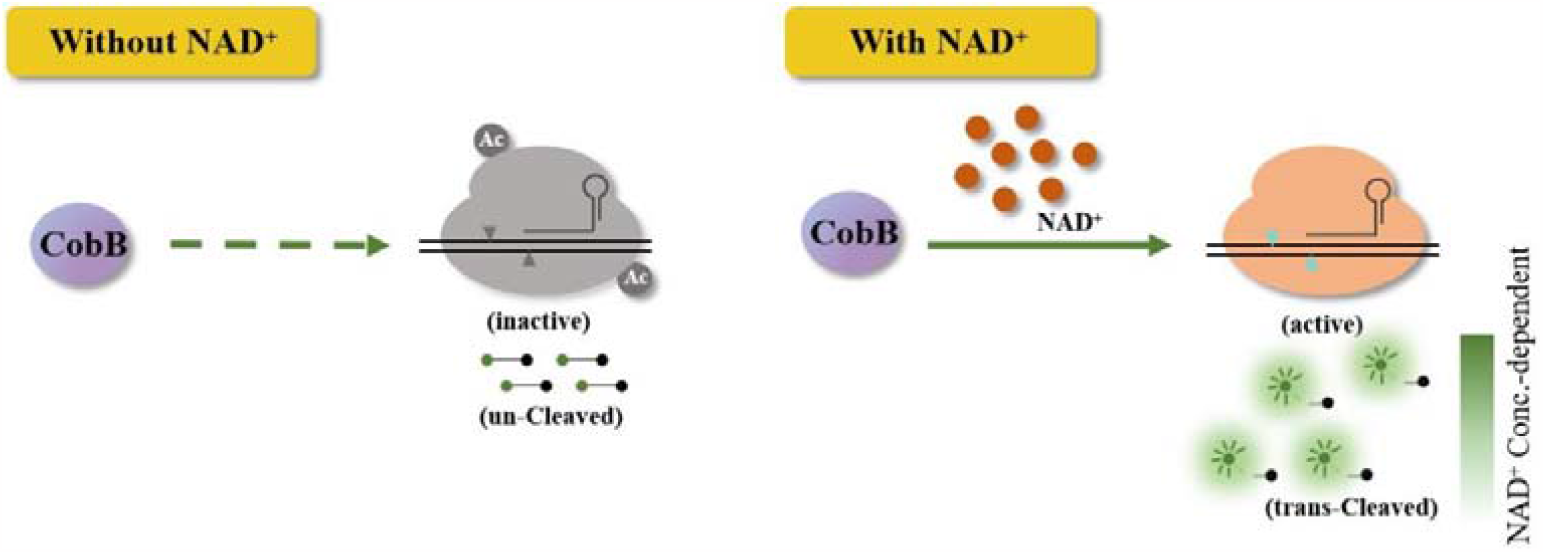
Schematic illustration of Cas12a-based CRISPR diagnostic system for NAD^+^ detection. Without NAD^+^, acetylated Cas12a losses the *trans*-cleavage activities and is unable to *trans*-cleave ssDNA reporters. When NAD^+^ is added, Cas12a in HOLMES(NAD) is activated, which results in the cleavage of reporters and illumination of fluorescence signals.

To test the hypothesis, we first need to produce acetylated Cas12a and establish the HOLMES(NAD) system. Usually, acetylation process could be performed by *in vitro* incubation of purified recombinant Cas12a together with AcrVA5 as previously described, where AcrVA5 efficiently acetylates the key lysine residue in Cas12a at the presence of acetyl-CoA (Kang et al. 2022). However, a further step is required to purify the acetylated Cas12a after the *in vitro* reaction, which procedure is time-consuming and could be harmful to both activity and stability of Cas12a (Fig. S1-3). Therefore, we here used an *in vivo* acetylation strategy by purification of acetylated LbCas12a from *Escherichia coli* BL21(DE3) co-expressing both LbCas12a and AcrVA5. Western blotting results showed that purified LbCas12a protein was efficiently acetylated (Fig. S4), and the acetylated LbCas12a (acLbCas12a) lost both *cis-* and *trans-*cleavage activities with target dsDNA (Fig. S5-7), which was consistent with our previous study (Kang et al. 2022) and demonstrated the practicability of the *in vivo* acetylation strategy. We further validated that acLbCas12a could be efficiently deacetylated by CobB (Fig. S8) and recovered its *cis-* and *trans-*cleavage activities when NAD^+^ cofactor was added. The restored *cis-* and *trans*-cleavage enzymatic activities were comparable to those of the control group using unacetylated Cas12a (Fig. S5-7). Noticeably, the reaction system with acLbCas12a, CobB, crRNA, target dsDNA and ssDNA FQ-reporters but without NAD^+^ is named the HOLMES(NAD) system.

### 3.2. Performance of HOLMES(NAD)

The HOLMES(NAD) stability was then analyzed, and no significant difference was found in fluorescence intensities after 20 times of repeated freeze-thaw treatment of the enzyme mixture of Cas12a and CobB (Fig. S9). When HOLMES(NAD) reactions were tested under temperatures ranging from 30 □ to 45 □, the most optimal temperature was determined to be 37 □ (Fig. S10). In addition, we found that HOLMES(NAD) was of both high repeatability (Table S5) and high precision (Table S6). Therefore, the above results clearly showed that HOLMES(NAD) could be triggered by NAD^+^ cofactor to illuminate fluorescence signals, demonstrating the system could be employed for stable and accurate NAD^+^ detection.

The limit of detection (LOD) of HOLMES(NAD) was further determined to be 22.5 nM, using serially diluted NAD^+^ solutions ranging from 1 mM to 7.8 nM. We found that the illuminated fluorescence signals gradually increased with the increase of the NAD^+^ concentration (Fig. 2A-B), and the value of the coefficient of determination (R^2^) was 0.997 between the fluorescence intensity and the NAD^+^ concentrations (from 1 mM to 31.3 nM) (Fig. 2C). The high R^2^ value not only reflected the linear relationship between added NAD^+^ concentration and the activated LbCas12a *trans*-cleavage activities but also indicated that HOLMES(NAD) could be employed for quantitative determination of NAD^+^ concentration.

**Fig 2.**
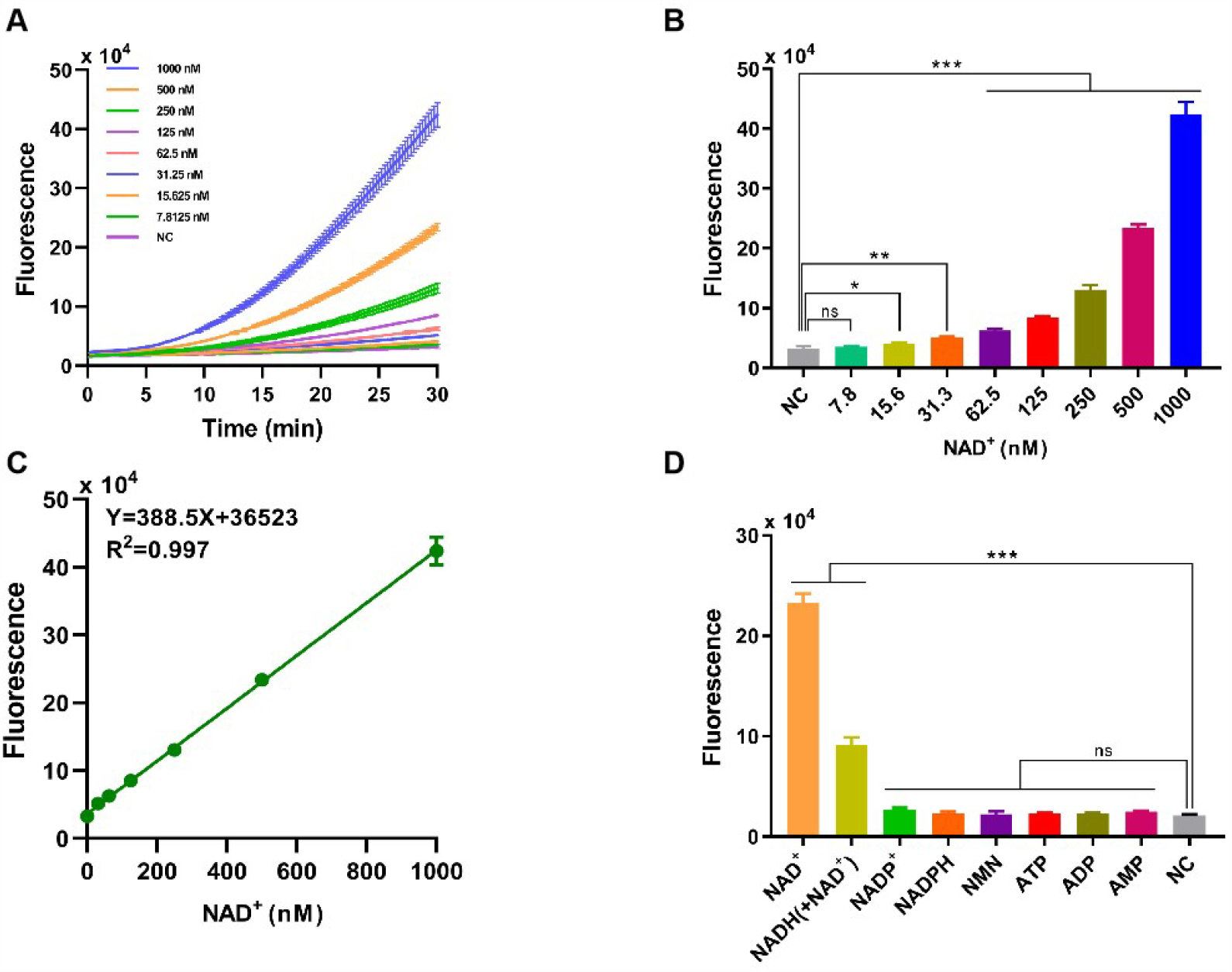
(A) Time-dependent fluorescence intensities of the HOLMES(NAD) in the presence of different concentrations of NAD^+^. (B) Fluorescence intensities of the HOLMES(NAD) with different concentrations of NAD^+^ added. Concentrations of NAD^+^ ranged from 7.8 nM to 1000 nM, and 30 min’s reaction was performed (n=3 technical replicates; two tailed Student’s t-test; ^*^ p<0.1; ^**^ p<0.01; ^***^ p<0.001; error bars represented the mean with SD). (C) Correlation between the fluorescence signals and the NAD^+^ concentrations. The solid line represents linear regression fitting to the experimental data. Three replicates were performed for each NAD^+^ concentration. (D) Fluorescence intensities of HOLMES(NAD) with various molecules added (n=3 technical replicates; two tailed Student’s t-test; ^***^ p<0.001; error bars represented the mean with SD).

The specificity of the HOLMES(NAD) system was then tested with the employment of various small molecules, including similar nucleotides and precursors for NAD^+^ biosynthesis. As expected, although some small molecules such as AMP, ADP, ATP, NMN, NADPH and NADP^+^ may share similar molecular structures with NAD^+^, they are not the cofactors of CobB and are theoretically unable to restore the *trans*-cleavage activities of acLbCas12a, resulting in no fluorescence signals output. However, to our great surprise, besides NAD^+^, HOLMES(NAD) also illuminated fluorescence signals when NADH was supplemented, which is inconsistent with the fact that NAD^+^ is the only cofactor of CobB (Fig. 2D). We therefore hypothesized that the commercially available NADH was contaminated with NAD^+^, which resulted in above unexpected signal output for NADH. To test this hypothesis, NADH was then measured by the mass spectrometry (MS) method, and the tested NADH solution was found to be a mixture of NADH and NAD^+^, which could therefore explain the above unexpected results for NADH. Moreover, the concentration of NAD^+^ in the NADH solution (500 nM) was 154.5 nM when measured by the quantitative MS method, which was consistent with the HOLMES(NAD) method, that is 140.8 nM (Fig. S11-14). Taken together, the above findings not only proved the high specificity of the HOLMES(NAD) method but also demonstrated its ability in target NAD^+^ quantitation.

### 3.3. Measurement of NAD^+^ levels in biological samples

Besides pure NAD^+^ solutions, we also verified HOLMES(NAD) system with extracts from real biology samples. Total NAD^+^ was first extracted from different cancer cell lines by heating at 95 □ and then neutralized before being detected. Using the HOLMES(NAD) method, the cellular NAD^+^ levels were identified to be approximately 200-600 pmol per million cells. The samples were also measured with other methods in parallel, and the results were consistent with those of the MS method and the MTT method (Fig. 3A). Moreover, the HOLMES(NAD) method was further used to quantitatively measure the changes of the cellular NAD^+^ concentrations, including 97H, HeLa and HCT116. After being treated with FK866, an inhibitor of the nicotinamide phosphoribosyltransferase, the cellular NAD^+^ levels dropped in all tested cell lines (Fig. 3B), which is consistent with the previous report (Pittelli et al. 2010).

**Fig 3.**
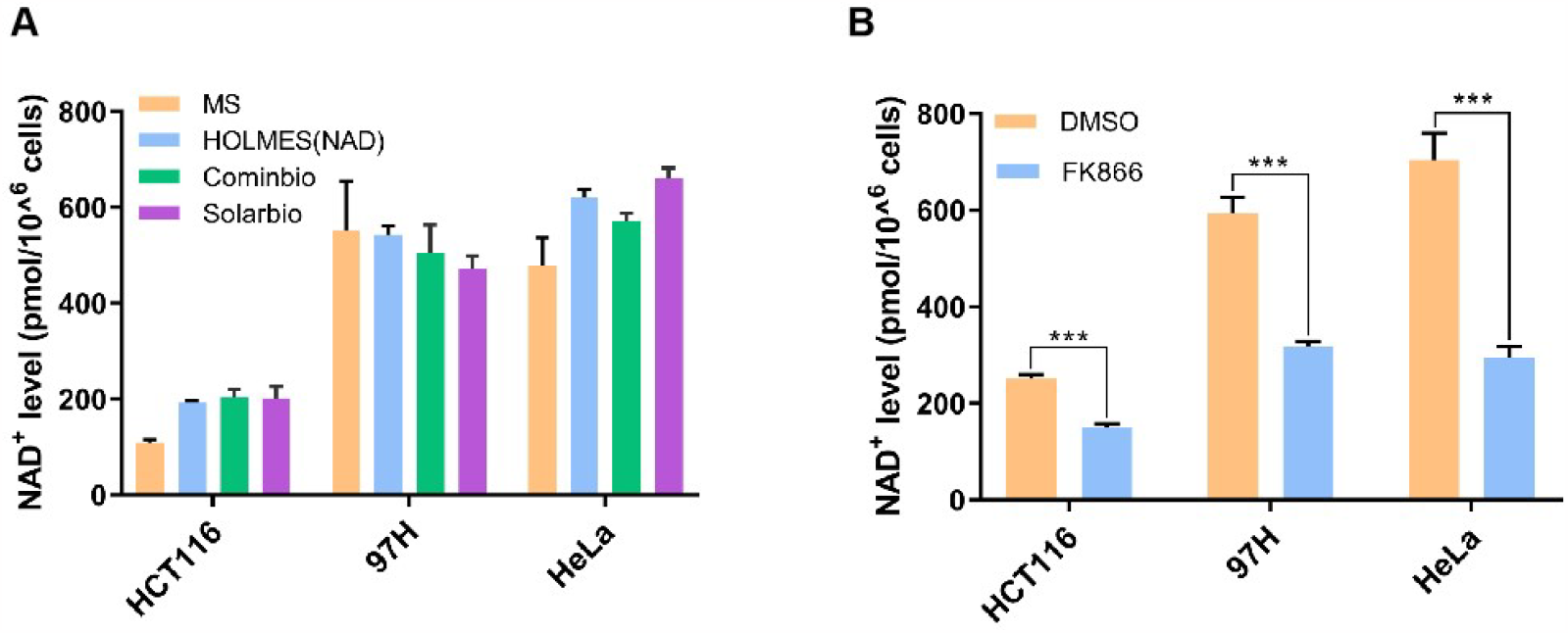
(A) Quantitative detection of the cellular NAD^+^ levels. Three cancer cell lines of HCT116, 97H, and HeLa were analyzed with three different methods, including the MS method, HOLMES(NAD), and the MTT-based NAD^+^ detection method. Two different commercially available MTT kits from Cominbio and Solarbio were used. (B) Quantitative detection of the changes of the cellular NAD^+^ levels after the treatment of FK866 in cell lines of HCT116, 97H and HeLa (n=3 technical replicates; two tailed Student’s t-test; ^***^ p<0.001; error bars represented the mean with SD).

## 4. Conclusion

In summary, we here create a CRISPR-Cas12a-based NAD^+^-specific diagnostic system, namely HOLMES(NAD). Through comparison of different NAD^+^ detection methods, we find that different methods show distinct advantages and disadvantages (Table S7). For example, although the MS method requires expensive instruments and professional operations, it can measure more than one target molecules in one test and allow for organic extraction that may better preserves molecules such as NADH and NADPH (Yamada et al. 2006). HOLMES(NAD) and other methods, by contrast, detect merely target NAD^+^ concentration, which could be less convenient when detection of multiple molecules is needed. Overall, HOLMES(NAD) shows apparent advantages in rapidness (half an hour), convenience, cost-effective, and accuracy in quantitative detection of NAD^+^ concentrations, which therefore has great potential in both scientific research and industry. Furthermore, this study expanded the application scenarios of the Cas12-based HOLMES diagnostic system from target nucleic acid detection to non-nucleic acid detection.

### CRediT authorship contribution statement

**Songkuan Zhuang:** Conceptualization, Methodology, Validation, Data Curation, Writing-Original Draft. **Tianshuai Hu:** Conceptualization, Validation. **Hongzhong Zhou:** Resources, Data Curation. **Shiping He:** Resources, Data Curation. **Jie Li:** Validation. **Dayong Gu:** Supervison, Writing-Review & Editing, Funding acquisition. **Yong Xu:** Supervision, Funding acquisition. **Yijian Chen:** Supervison, Writing-Review & Editing, Funding acquisition. **Jin Wang:** Conceptualization, Supervison, Methodology, Writing-Review & Editing, Funding acquisition.

### Data availability

Data will be made available on request.

## Acknowledgment

All authors made contributions to the manuscript and have approved the final version of the manuscript. This study was financially supported by the National Natural Science Foundation of China (31922046), the National Key Research and Development Program of China (2022YFC2302700), the Guangdong Science and Technology Foundation (2021A1515220084 and 2022B1111020001), Shenzhen Science and Technology Program (ZDSYS20210623092001003, GJHZ20200731095604013, JSGG20220301090003004, 201906133000069, SGLH20180625171602058 and JCYJ20200109120205924) and Shanghai Science and Technology Innovation action plan of the 2022 in agricultural field (22N31900200).

## Appendix A. Supplementary data

Supplementary data to this article can be found online at xx.

