## Supplementary material for "CRISPR-HOLMES-based NAD^+^ detection": Supplementary material.docx

**Running title:** NAD^+^ detection by HOLMES

Songkuan Zhuang, Tianshuai Hu, Hongzhong Zhou, Shiping He, Jie Li, Yuehui Zhang, Dayong Gu, Yong Xu*, Yijian Chen^*^ and Jin Wang^*^

^#^These authors contributed equally to this work.

^*^Correspondence should be addressed to Yong Xu, Yijian Chen and Jin Wang.

**CONTENTS**

**Table S1.** Oligonucleotides used for plasmids construction in this study.

**Table S2.** Oligonucleotides used for preparation of LbCas12a targets in this study.

**Table S3.** The crRNA and FQ-reporter used in this study.

**Table S4.** Similar nucleotides and biosynthetic precursors for NAD^+^.

**Table S5.** Repeatability test of HOLMES(NAD) detection methods.

**Table S6.** Precision test of HOLMES(NAD) detection methods.

**Table S7.** Comparison of different NAD^+^ detection methods.

**Fig. S1.** Western blotting analysis of the lysine acetylation levels using acLbCas12a produced by both *in vitro* and *in vivo* approaches.

**Fig. S2.** SDS-PAGE analysis of purified recombinant CobB protein.

**Fig. S3.** Time-dependent fluorescence intensities of the Cas12a *trans*-cleavage experiments.

**Fig. S4.** Western blotting analysis of the acetylated LbCas12a (acLbCas12a).

**Fig. S5.** *Cis*-cleavage assay with acLbCas12a.

**Fig. S6.** Time-dependent fluorescence intensities of the acLbCas12a *trans*-cleavage experiment.

**Fig. S7.** *Trans*-cleavage assay with acLbCas12a.

**Fig. S8.** WB analysis of the acetylation status of acLbCas12a.

**Fig. S9.** The effect of freeze-thaw treatment on the stability of the proteins in HOLMES(NAD) system.

**Fig. S10.** Analysis of the applicable temperature range for HOLMES(NAD).

**Fig. S11.** Analysis of the composition of the NAD^+^ standards with mass spectrometry (MS).

**Fig. S12.** Analysis of the composition of the NADH standards with mass spectrometry (MS).

**Fig. S13.** Quantitative analysis of the NAD^+^ and NADH contents in the NADH standards by mass spectrometry.

**Fig. S14.** Calculation of the NAD^+^ contents in the NADH standards.

**Table S1. Oligonucleotides used for plasmids construction in this study.**

| **Oligo names** | **Sequences (5'-3')** |
| --- | --- |
| LbCas12a-F | cgcGGATCCcatgctgaagaacgtgggcatcga |
| LbCas12a-R | cgcGTCGACtcagtgtttcacgctggtctgagcata |

**Table S2. Oligonucleotides used for preparation of LbCas12a targets in this study.**

| **Oligo names** | **Sequences (5'-3')** |
| --- | --- |
| T1-50 nt-F | gttgtaaaacgacggccagttttgttatcgcaactttctactgaattcgg |
| T1-50 nt-R | ccgaattcagtagaaagttgcgataacaaaactggccgtcgttttacaac |
| M13-F | TGTAAAACGACGGCCAGT |
| M13-R-FAM | FAM-CAGGAAACAGCTATGACC |

**Table S3. The crRNA and FQ-reporter used in this study.**

| **Oligo names** | **Sequences (5'-3')** |
| --- | --- |
| crRNA | AAUUUCUACUCUUGUAGAUUUAUCGCAACUUUCUACUGAAUU |
| FQ-reporter | 5’-FAM-CCCCCCCC-BHQ1-3’ |

**Table S4. Similar nucleotides and biosynthetic precursors for NAD^+^.**

| **Name** | **Provider** | **Cat. Nos.** |
| --- | --- | --- |
| NAD^+^ | Sigma | N7004-250MG |
| NADH | MCE | HY-F0001 |
| NADP^+^ | MERCK | 24292-60-2 |
| NADPH | Sigma | N1630-250MG |
| NMN | RHAWN | R019041-100MG |
| ATP | Sigma | A6419-1G |
| ADP | YONGQI | 20398-34-9 |
| AMP | YONGQI | 4578-31-8 |

**Table S5. Repeatability test of HOLMES(NAD) detection methods.**

| **NAD^+^ conc. (nM)** | **1**  **(Fluorescence)** | **2**  **(Fluorescence)** | **3**  **(Fluorescence)** | **Mean**  **(Fluorescence)** | **RSD**  **(%)** |
| --- | --- | --- | --- | --- | --- |
| 1,000 | 430406.063 | 464732.469 | 449093.844 | 448077.458 | 3.84 |
| 600 | 277618.563 | 257968.734 | 263600.938 | 266396.078 | 3.80 |
| 150 | 85089.156 | 95409.117 | 92321.094 | 90939.789 | 5.82 |

**Table S6. Precision test of HOLMES(NAD) detection methods.**

| **NAD^+^ conc. (nM)** | **1**  **(nM)** | **2**  **(nM)** | **3**  **(nM)** | **4**  **(nM)** | **5**  **(nM)** | **Mean**  **(nM)** | **RSD**  **(%)** | **Measured**  **/True values** |
| --- | --- | --- | --- | --- | --- | --- | --- | --- |
| 1,000 | 1055.263 | 1070.972 | 1103.856 | 1083.544 | 1081.335 | 1078.994 | 1.65 | 1.079 |
| 500 | 517.028 | 489.253 | 532.828 | 509.723 | 495.917 | 508.950 | 3.39 | 1.018 |
| 100 | 103.955 | 80.211 | 89.669 | 93.992 | 99.005 | 93.367 | 8.72 | 0.934 |

**Table S7. Comparison of different NAD^+^ detection methods.**

|  | Sensitivity | Specificity | Accurate | Labor  intensive | Expensive equipment | Time/min | References |
| --- | --- | --- | --- | --- | --- | --- | --- |
| Enzymatic assay | 8.3 nM | YES | YES | YES | NO | 50 | [1] |
| HPLC-UV | 1 μM | YES | YES | YES | YES | 40 | [2] |
| LC/MS/MS | 10 nM | YES | YES | YES | YES | 35 | [3] |
| NAD-CPLuc^2^ | 20 nM | NO | YES | NO | NO | 10 | [4] |
| HOLMES(NAD) | 22.5 nM | YES | YES | NO | NO | 30 | This work |

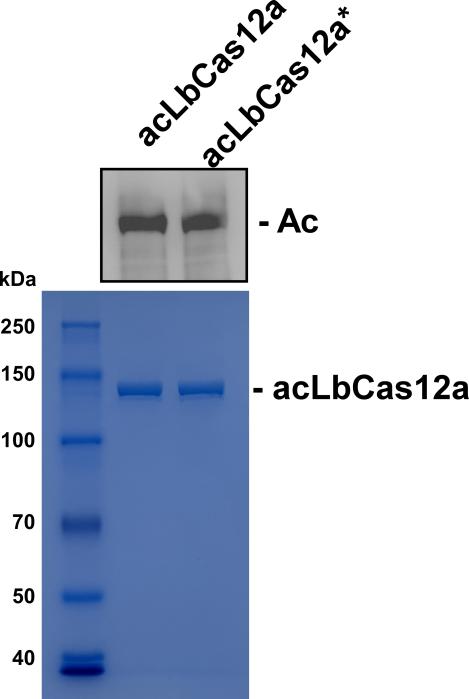

**Fig. S1. Western blotting analysis of the lysine acetylation levels using acLbCas12a produced by both *in vitro* and *in vivo* approaches.** The acLbCas12a proteins were obtained from *E. coli* co-expressing both LbCas12a and AcrVA5, designated as the *in vivo* method. The acLbCas12a* proteins were obtained by AcrVA5-mediated *in vitro* acetylation treatment of LbCas12a protein.

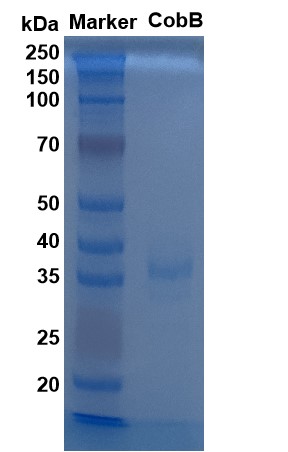

**Fig. S2. SDS-PAGE analysis of purified recombinant CobB protein.**

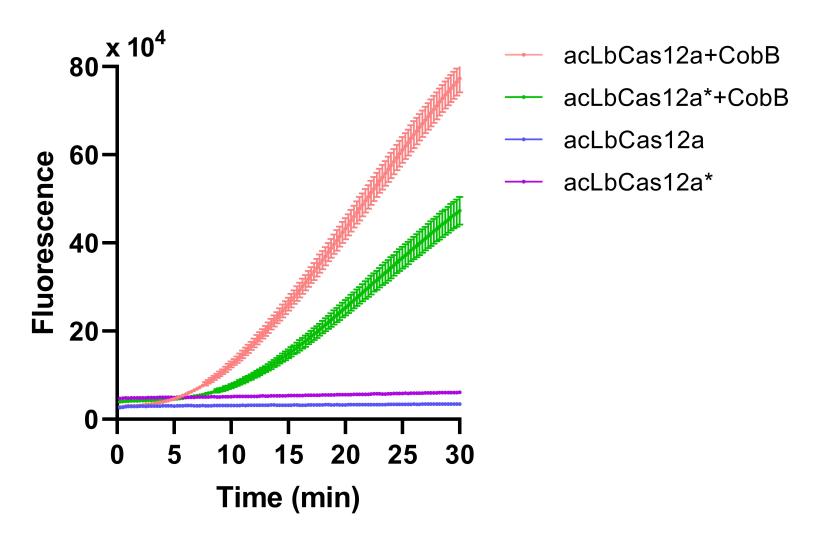

**Fig. S3. Time-dependent fluorescence intensities of the Cas12a *trans*-cleavage experiments.** Assays were performed with different acLbCas12a obtained from either *in vivo* or *in vitro* (*) methods. acLbCas12a was treated with or without the addition of CobB deacetylation components.

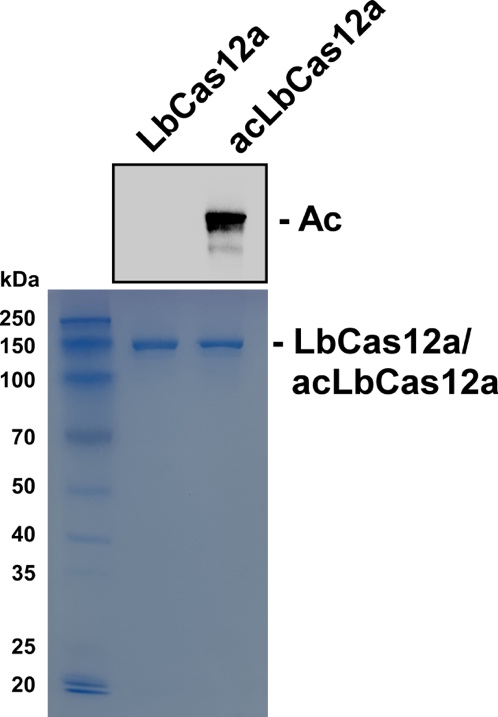

**Fig. S4.** **Western blotting analysis of the acetylated LbCas12a (acLbCas12a).** Both LbCas12a and acLbCas12a were analyzed by SDS-PAGE, followed by WB analysis.

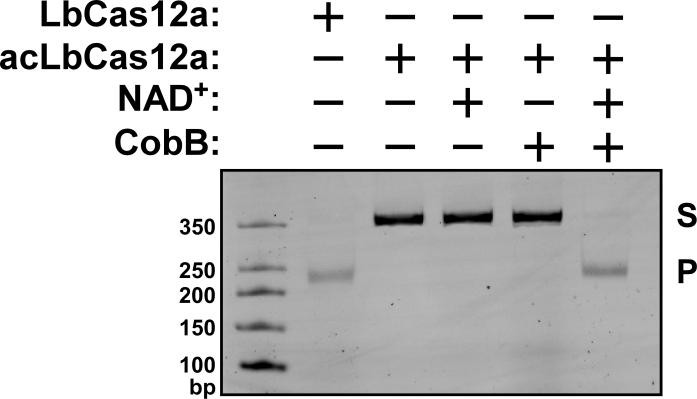

**Fig. S5. *Cis*-cleavage assay with acLbCas12a.** acLbCas12a was treated with different conditions with or without CobB and NAD^+^. S, dsDNA substrate; P, Cas12a *cis*-cleaved products.

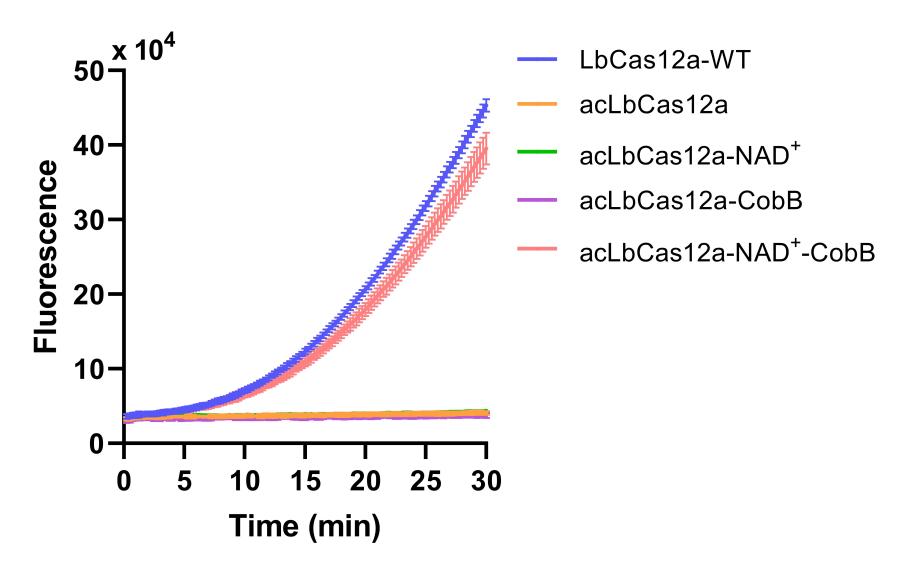

**Fig. S6. Time-dependent fluorescence intensities of the acLbCas12a *trans*-cleavage experiment.** acLbCas12a was treated in conditions with or without the addition of CobB and NAD^+^.

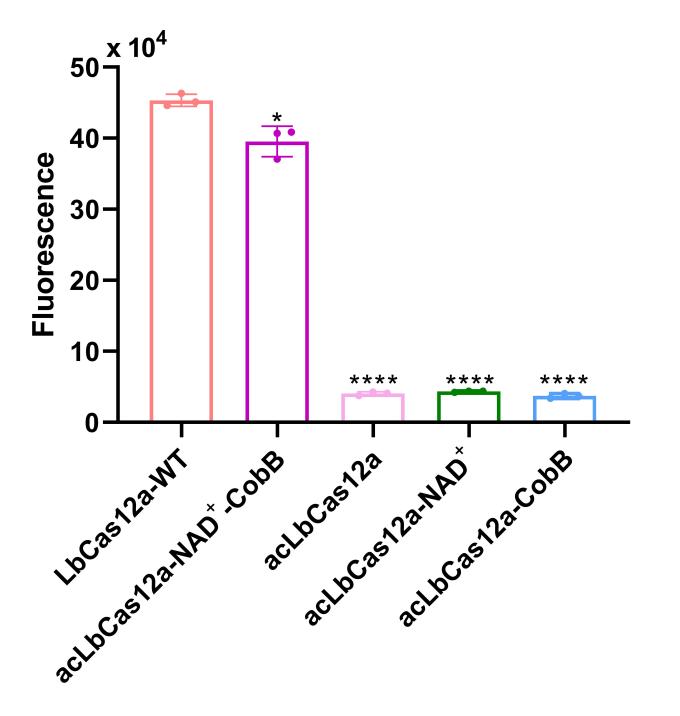

**Fig. S7. *Trans*-cleavage assay with acLbCas12a.** Assay was performed with acLbCas12a treated with or without the addition of CobB and NAD^+^. The presented fluorescence intensities were obtained by incubating the reaction at 37 ℃ for 30 min (n=3 technical replicates; two tailed Student’s t-test; * p<0.1; **** p<0.0001; error bars represent the mean with SD).

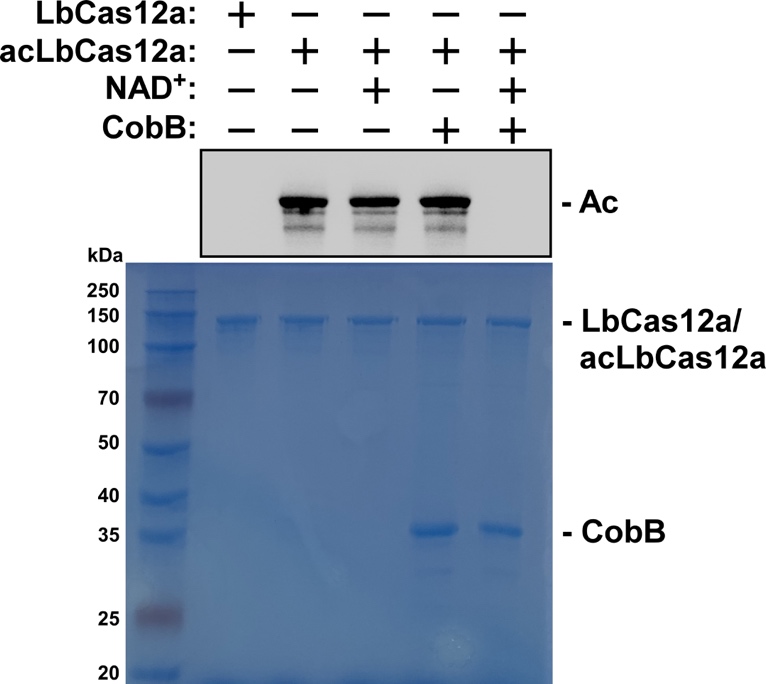

**Fig. S8. WB analysis of the acetylation status of acLbCas12a.** acLbCas12a was treated by CobB-mediated deacetylation in conditions with or without the addition of CobB and NAD^+^. α-Kac, pan anti-acetyl lysine antibody (PTM Bio, Cat# PTM-001).

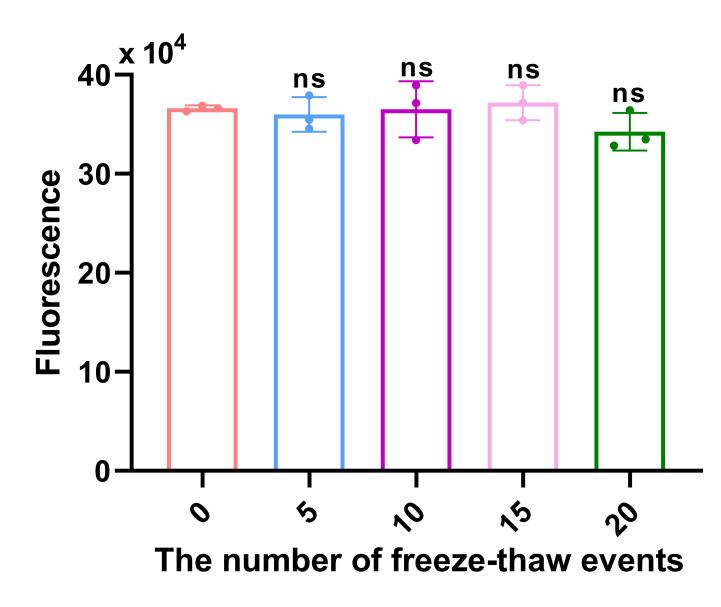

**Fig. S9. The effect of freeze-thaw treatment on the stability of the proteins in HOLMES(NAD) system.** The acLbCas12a/CobB protein mixture was first frozen at -80 ℃ and then thawed at 37 ℃. After repeated freeze-thaw treatment, the protein mixture was used for HOLMES(NAD) analysis. The fluorescence intensities were obtained through incubating the reaction at 37 ℃ for 30 min (n=3 technical replicates; two tailed Student’s t-test; error bars represent the mean with SD).

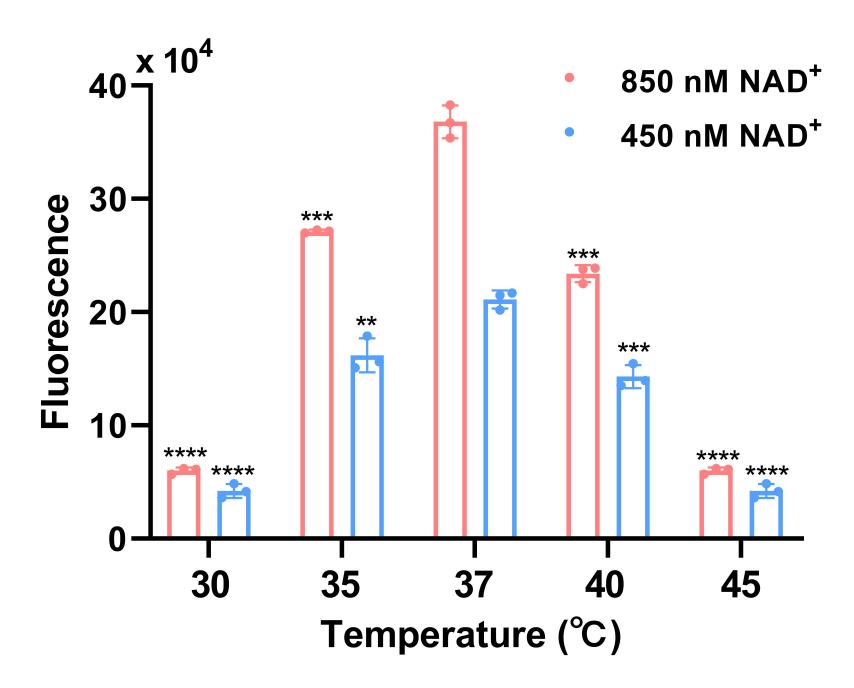

**Fig. S10. Analysis of the applicable temperature range for HOLMES(NAD).** HOLMES(NAD) reaction was performed at designated temperatures using two different NAD^+^ concentrations of both 850 nM and 450 nM. The fluorescence intensities were obtained through incubating the reaction for 30 min (n=3 technical replicates; two tailed Student’s t-test; ** p<0.01; *** p<0.001; **** p<0.0001; error bars represent the mean with SD).

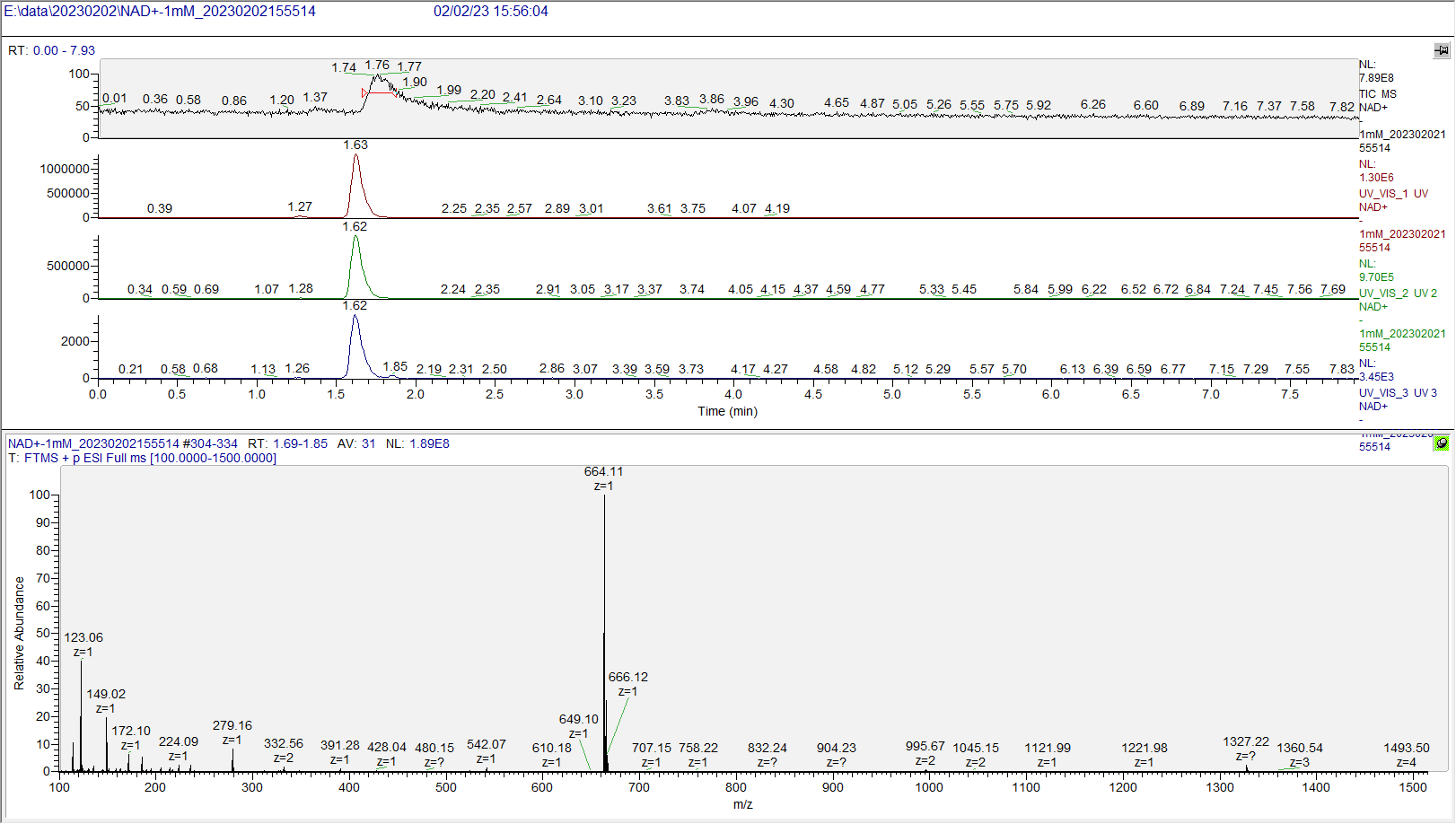

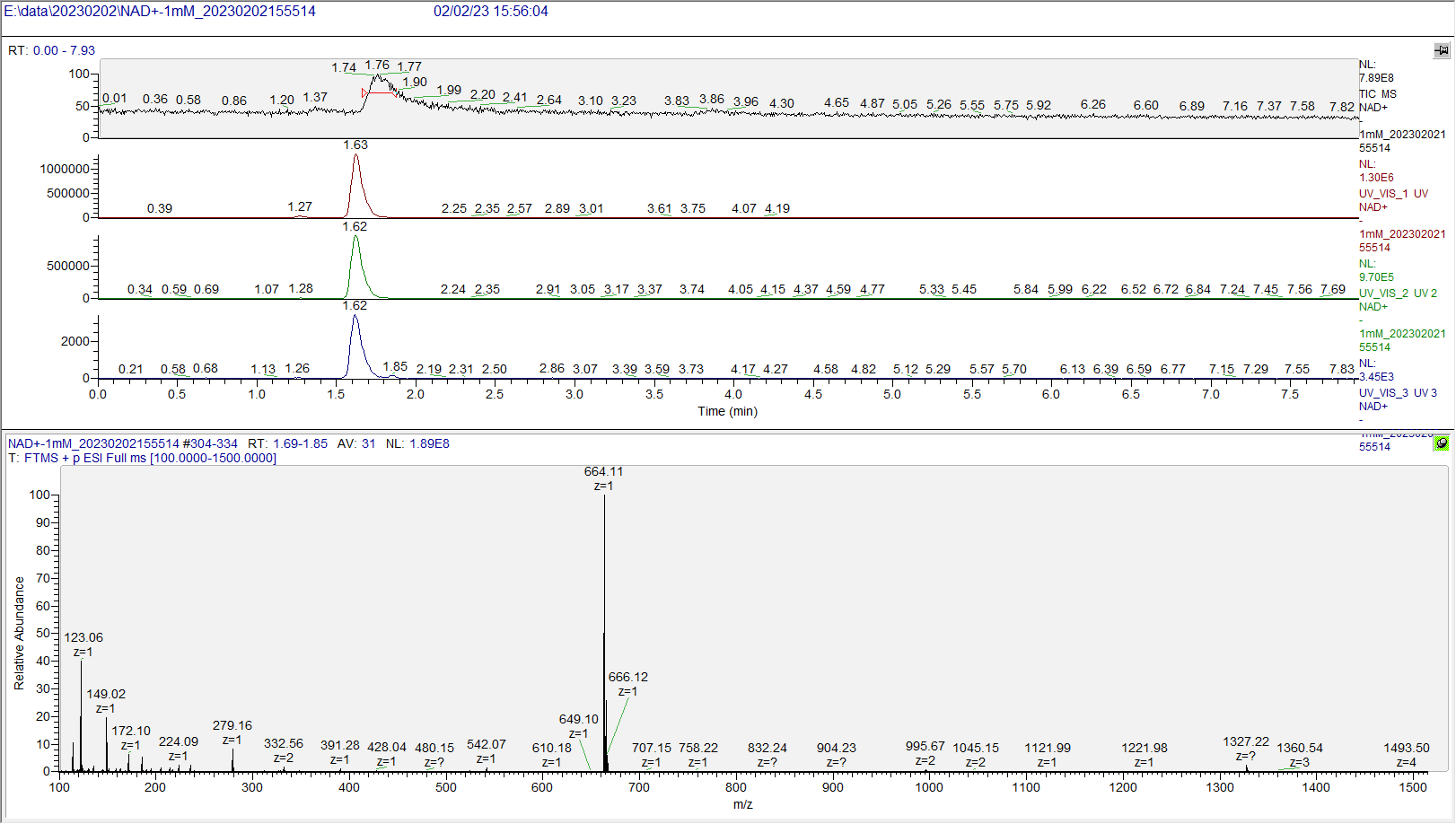

**Fig. S11.** **Analysis of the composition of the NAD^+^ standards with mass spectrometry (MS).** Main component of the MS signals collected at retention time from 1.69 to 1.85 minutes was NAD^+^.

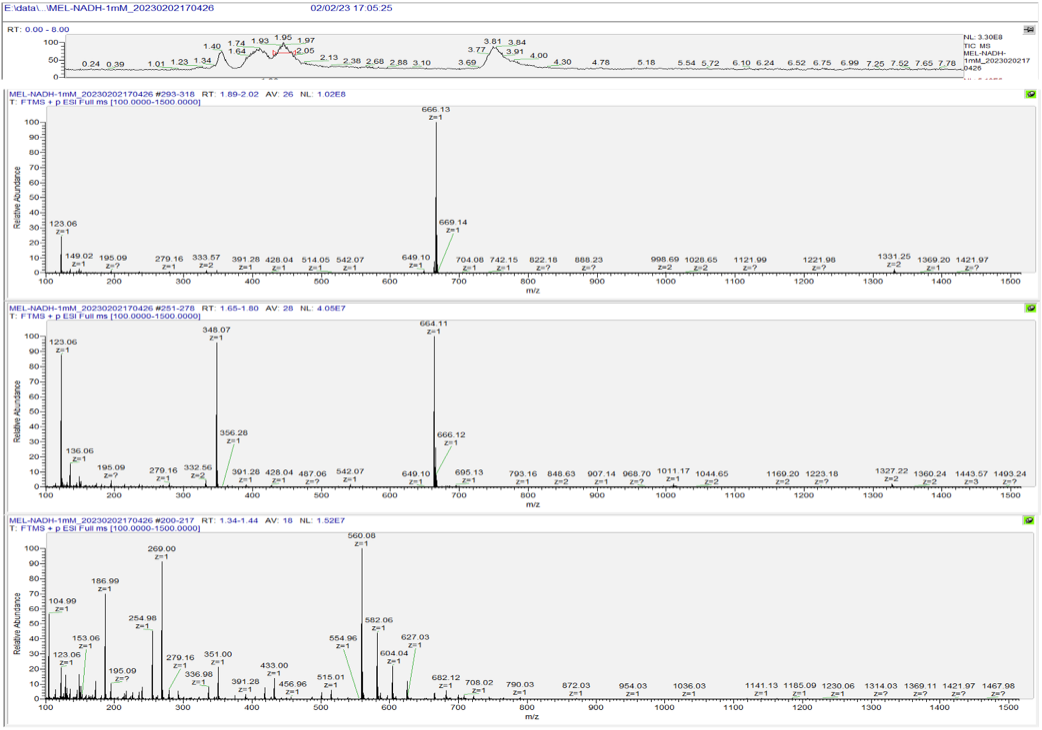

**Fig. S12. Analysis of the composition of the NADH standards with mass spectrometry (MS).** Main component of the MS signals collected at retention time from 1.89 to 2.02 minutes was NADH, while the component from 1.65 to 1.80 minutes was NAD^+^. Components with retention time from 1.34 to 1.44 minutes were contaminants.

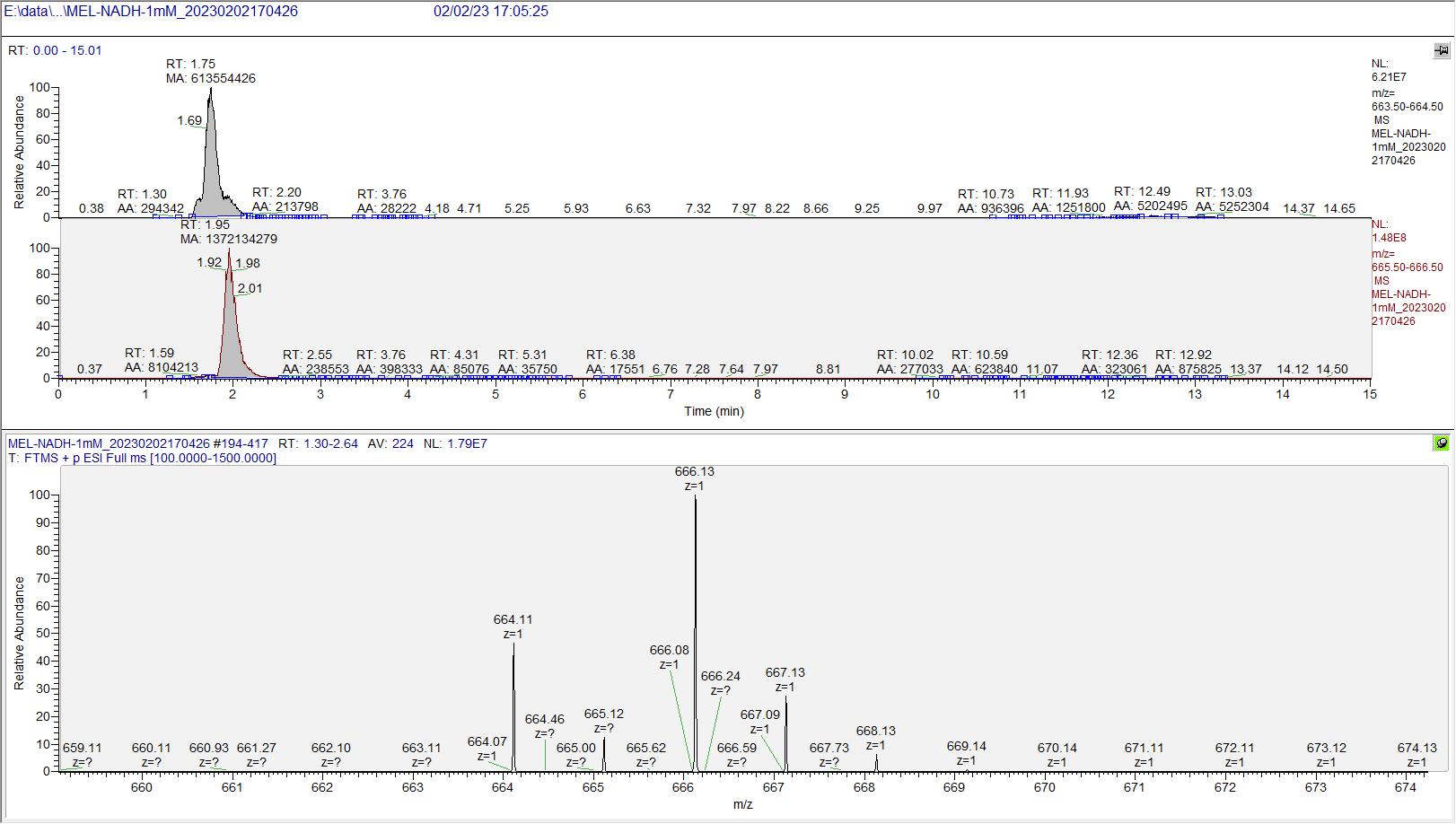

**Fig. S13. Quantitative analysis of the NAD^+^ and NADH contents in the NADH standards by mass spectrometry.** The NAD^+^ peak area was 613,554,426, and the NADH peak area was 1,372,134,279.

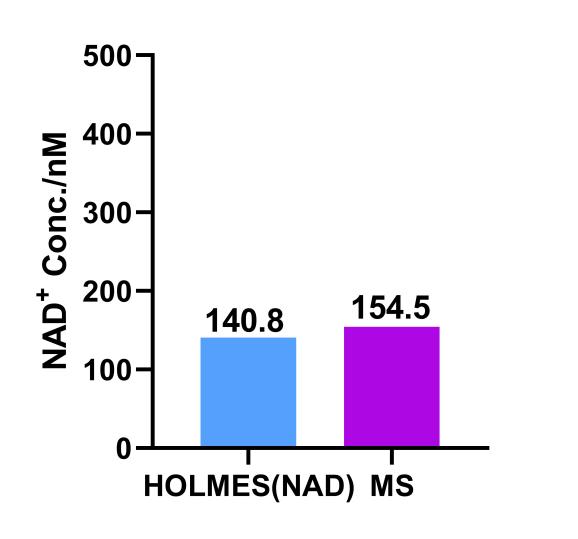

**Fig. S14. Calculation of the NAD^+^ contents in the NADH standards.** The contents were analyzed with both the HOLMES(NAD) method and the MS method. For the HOLMES(NAD) method, fluorescence intensities of the NADH standard were converted to the NAD^+^ concentration using the formula from the standard curve, which is Y= 388.5X +36523. While for the MS method, the NAD^+^ content was determined using the following formula the concentration of [NADH]*(peak areas [NAD^+^]/peak areas ([NAD^+^]+[NADH])).
